# iATPSnFR2: a high dynamic range fluorescent sensor for monitoring intracellular ATP

**DOI:** 10.1101/2023.08.24.554624

**Authors:** Jonathan S. Marvin, Alexandros C. Kokotos, Mukesh Kumar, Camila Pulido, Ariana N. Tkachuk, Jocelyn Shuxin Yao, Timothy A. Brown, Timothy A. Ryan

## Abstract

We developed a significantly improved genetically encoded quantitative adenosine triphosphate (ATP) sensor to provide real-time dynamics of ATP levels in subcellular compartments. iATPSnFR2 is a variant of iATPSnFR1, a previously developed sensor that has circularly permuted super-folder GFP inserted between the ATP-binding helices of the *ε*-subunit of a bacterial F_0_-F_1_ ATPase. Optimizing the linkers joining the two domains resulted in a ∼ 5-6 fold improvement in the dynamic range compared to the previous generation sensor, with excellent discrimination against other analytes and affinity variants varying from 4 μM to 500 μM. A chimeric version of this sensor fused to either the HaloTag protein or a suitably spectrally separated fluorescent protein, provides a ratiometric readout allowing comparisons of ATP across cellular regions. Subcellular targeting of the sensor to nerve terminals reveals previously uncharacterized single synapse metabolic signatures, while targeting to the mitochondrial matrix allowed direct quantitative probing of oxidative phosphorylation dynamics.

**Significance Statement:** Adenosine triphosphate (ATP) is a key metabolite necessary for cellular life. Here we develop a next-generation genetically encoded ratiometric fluorescent ATP sensor that allows subcellular tracking of ATP levels in living cells. The large dynamic range makes it possible to follow the dynamics of this metabolite across cells and subcellular regions under different metabolic stressors. We expect that iATPSnFR2 will provide researchers with exciting new opportunities to study ATP dynamics with temporal and spatial resolution that has, until now, been unavailable.

## Introduction

ATP is a critical biochemical currency. Its hydrolysis to ADP provides the free energy necessary to drive numerous physiological processes. The importance of ATP is evident by the plethora of routes that exist to convert the energy stored in the form of combustible hydrocarbons into ATP. The essentials of glycolysis and oxidative phosphorylation have been established for over sixty years, having been successfully dissected through *in vitro* enzymology and biochemistry. One trade-off with this type of reductionism is that it is often hard to recapitulate the physiological milieu *in vitro*, and therefore the nuances of how molecular pathways are regulated in real time in specific subcellular locations are missed. Fluorescent biosensors are designed to fill the knowledge gap between *in vitro* reconstitution biochemistry and cellular physiology as they can reveal subcellular dynamics of different metabolites or ions with high temporal and spatial resolution in living cells and tissues. Successful deployment of a fluorescent biosensor requires that the affinity of the sensor matches the physiological concentration of its cognate analyte and that the sensor discriminates against chemically related species. The signal-to-noise ratio is determined by the ratio of the sensor’s fluorescence in the bound and unbound states and sets the limit of what magnitude changes in the analyte concentration can be detected in the face of other sources of fluctuation and therefore the spatial scale over which the signal must be averaged. Several promising strategies have emerged in the last ten years to develop a genetically encoded sensor for ATP.

An intensity-based ATP Sensing Fluorescent Reporter, iATPSnFR(1), was developed by inserting circularly permuted superfolder GFP between the two ATP-binding helices of the epsilon subunit of the F_0_-F_1_ ATPase of thermostable *Bacillus subtilis* PS3, which undergoes a large conformational change upon binding ATP(2). Other research groups have made similarly conceived sensors, albeit with different topologies, ATP affinities, and dynamic ranges. These include ChemoG-ATP_SiR_(3), MaLion(4), and QUEEN(5), which, like iATPSnFR have a common ancestor in “ATeam”, a FRET based sensor in which CFP and YFP were placed at the N- and C-termini of this bacterial *ε* subunit of F_0_-F_1_ ATPase(6), and whose FRET efficiency improves as this subunit changes conformation up binding ATP. These cytosolic ATP sensors have been reviewed recently(7). Unfortunately, each of these sensors has been lacking in some respect. QUEEN detects changes in ATP by reporting different excitation wavelengths, which is not practically useful for most imaging experiments, especially for preparations other than monolayers. MaLion is available as either a green and or red fluorescent sensor and has almost 5-fold increase in fluorescence upon saturation with ATP, but loses dynamic range at higher temperatures. The most recently developed ATP sensor, ChemoG-ATP_SiR_, is an improved chemogenetic FRET sensor with 10-fold change in fluorescence ratio(3). ChemoG-ATP_SiR_ has the necessary large dynamic range, and has an affinity of about 2 mM, which is in the physiological range for many cells. It also is available in different colors, depending on the FRET acceptor used. However, the use of two fluorophores occupies spectral bandwidth that might limit its application to multiplex imaging. Other ATP sensors based on the bacterial ATP regulatory domain GlnK1, including Perceval(8) and PercevalHR(9), have similar affinities for ATP and ADP, and thus provide a measure of the ratio of ATP:ADP more than the concentration of ATP alone. Finally, firefly luciferase can be used to measure the concentration of ATP, as there is a direct relationship between ATP consumption and light production. This approach is obviously limited in its spatial and temporal resolution, but was successfully adapted to observe ATP consumption at nerve terminals in primary dissociated neurons(10). Even so, that sensor is difficult to deploy due to the low photon flux and limited applicability to modern optical sectioning.

Here we report the improvement of iATPSnFR1(1). iATPSnFR2 has much higher dynamic range (ΔF/F ∼ 12), is available as three different affinity variants, and is fused to spectrally separable fluorescent tags (HaloTag with synthetic far-red fluorophores or mIRFP670nano3) thus providing an approach to normalize the signal to the expression level of the sensor and allowing quantitative comparisons across individual cells or subcellular locations. We show that iATPSnFR2 can provide detailed measurements of the variations in resting ATP values across synapses as well as the kinetics of ATP changes during metabolic perturbations at the cytosolic, single synapse, and single mitochondrial levels. These data show for the first time that individual synapses behave as semi-independent metabolic units, as during metabolic stress, the kinetics of ATP depletion varied significantly even within the same axon.

## Results

### Sensor design and *in vitro* characterization

To improve upon iATPSnFR, we first re-evaluated the composition of the linkers connecting the ATP-binding domain and the cpSFGFP (Fig. 1a). This was done with the inclusion of an ATP-affinity boosting mutation, A95K, as we were also aiming to create a high-affinity ATP sensor that would be useful for the detection of sub-micromolar amounts of extracellular ATP. To increase ΔF/F we screened thousands of variants of the sensor in bacterial lysate, as described previously(1), but expanded the regions mutated to cover a larger number of residues. The sensor with the highest maximum ΔF/F (saturated vs unbound) has the residues comprising “linker 1” as VLVG, where the residues QD adjacent to the ATP-binding domain are changed to VL, and the residues SH adjacent to cpSFGFP are changed to VG. Additional residues ICV were placed in “linker 2” between the end of cpSFGFP and residue 110 of the ATP-binding domain. This sensor (L1-VLVG, L2-ICV, A95K) has an affinity for ATP of ∼16 uM.

**Figure 1.**
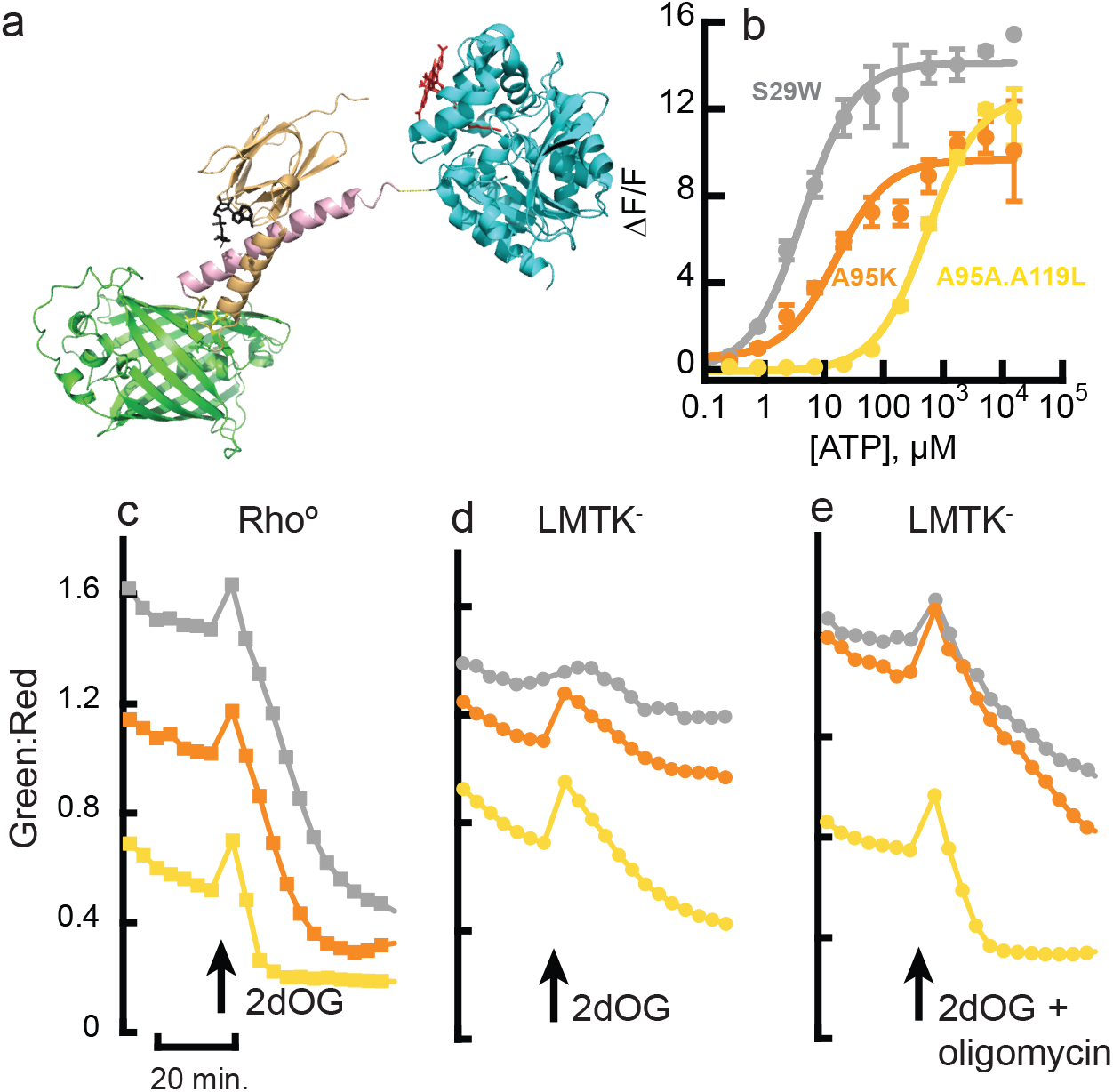
Design and characterization of iATPSnFR2 *in vitro* and in fibroblasts. a) Artistic rendering of iATPSnFR2.HaloTag made with PyMol. The N-terminal fragment (residues 1-109) of the epsilon subunit of ATPase (orange, based on 2E5Y.PDB) is fused to cpSFGFP (green). The C-terminus of cpSFGFP is fused to the residual ATP-binding helix (residues 110-129) of the epsilon subunit (pink), which itself is fused to HaloTag (cyan). “Linkers” are shown as sticks in yellow. ATP is shown as sticks in black. HaloTag fluorophore is shown as sticks in red. b) iATPSnFR2.HaloTag-JFX650 affinity variants titrated with ATP. Grey, S29W.A95K (4 μM); orange, A95K (16 μM), yellow, A95A.A119L (530 μM). c-e) Ratio of green (iATPSnFR2) to red (HaloTag-JFX650) fluorescence in fibroblasts transfected with three affinity variants of iATPSnFR2-HaloTag. Grey, S29W.A95K (high affinity); orange, A95K (medium affinity); yellow, A95A.A119L (low affinity). Cells were incubated for 21 minutes in buffer containing 10 mM glucose, and then switched to buffer containing 10 mM 2-deoxyglucose. c) Rhoº cells. d) Parental LMTK cell line. e) Parental cell line treated with both 2-deoxyglucose and oligomycin.

To tune the affinity of the sensor into the range needed for measuring cytosolic ATP in neurons (estimated to be ∼1.4 mM(10)), we reverted A95K back to alanine, which also reduced ΔF/F. We then screened residues near the ATP-binding site for variants that had a high ΔF/F upon saturation with ATP and had a *K*_d_ near 1.4 mM. The substitution A119L satisfied those two criteria and was carried forward as iATPSnFR2 (L1-VLVG, L2-ICV, A95A, A119L). In parallel, we screened other amino acid positions in the epsilon subunit in the context of A95K for variants that bound ATP more tightly. We found that the double mutant S29W.A95K has an affinity for ATP of ∼ 4 μM, giving us a trio of sensors with similar ΔF/F and varying affinities: S29W.A95K (4 μM), A95K (16 μM), A95A.A119L (530 μM). (Fig. 1b and Supp. Fig 1a).

Like GCaMP and other cpGFP-based sensors, the excitation peaks at 405 nm and 485 nm are shifted relative to each other upon saturation with ligand, providing an isosbestic point at 436 nm (Supp. Fig. 1b,c), which provides an ATP-independent signal that may be useful for normalization of focus and movement artefacts. However, since imaging with shorter wavelength excitation light is often detrimental to live cells, we also developed a version of the sensor that include a C-terminal fusion proteins to act as a normalization reference. A particularly useful C-terminal fusion protein is HaloTag, which can serve as a conjugation partner for synthetic fluorophores(11). There are many synthetic fluorophores available for conjugation to HaloTag, and for the present study we primarily used JF635 or JFX650 (12).

The iATPSnFR2.HaloTag-JFX650 was the primary protein used for *in vitro* characterization. The inclusion of HaloTag affects affinity, and also appears to increase the shelf-life of the purified sensor. iATPSnFR2.A95A.A119L.HaloTag-JFX650 has a ΔF/F of ∼12 and *K*_d_ for ATP of 500 μM (Fig 1b), when taken at the maximum point of excitation/emission sensitivity. It has low affinity (5.3 mM) and ΔF/F (∼6) for ADP (Fig 1b, Supp. Fig 1d). This version of the sensor has minimal, if any, affinity for AMP. It also changes fluorescence in response to GTP and CTP, but not TTP (Supp Fig. 1e). The affinity for these nucleoside triphosphates is low enough that their presence in cells should not affect sensor performance, given that reported concentrations of these compounds are about 0.5 mM (GTP) and 0.3 mM (CTP)(13). Reports of other ATP sensors based on the epsilon subunit of ATPase do not mention the affinity of those sensors for CTP or TTP. MaLion (4) and ChemoG-ATOP_SiR_(3) show no change in fluorescence with GTP. Finally, iATPSnFR2 is not affected by high concentrations of inorganic phosphate (Supp Fig. 1f).

MaLion has a markedly lower ΔF/F at 37°C than at 25°C (Supp. Fig. S4 of ref(4)), and ChemoG-ATP_SiR_ is also sensitive to temperature, but less so (Ext. Data Fig. 5d of ref(3)). In contrast, iATPSnFR2 has low temperature sensitivity, with almost identical ΔF/F and *K*_d_ for ATP at 37°C and 25°C (Supp Fig. 1g). Moreover, in contrast to MaLion, the rate of fluorescence change for all three sensors appears to be an order of magnitude faster for iATPSnFR2 (Supp Fig 1h) than the published values for MaLaion (Supp. Fig. S3 of ref.(4)). However, we did not do a head-to-head comparison of the two sensors, so they might in fact react at similar rates under similar conditions. Regardless, the A95K.S29W and A95K sensors bind ATP faster than the A95A.A119L variant and the rates of fluorescence increase do not follow a single exponential function (a phenomenon observed with other cpGFP-based sensors(14)), indicating a mechanism more complicated than concerted binding and increased fluorescence. Still, the sensors reach their maximum fluorescence within 1 to 2 seconds.

Finally, since deviation from metabolic homeostasis might result in changes in the pH of the cytosol, we characterized the pH dependence of iATPSnFR2. cpSFGFP has a pKa of about 7.2 and, as expected, does not respond to ATP (Supp Fig 1i, green lines). The ATP-bound form of the sensor (Supp Fig 1j, black solid line) shows a similar pH profile, but the ligand-free sensor has a lower pKa, closer to about 6.0 (Supp Fig 1h, black dashed line).

### Validation in cell lines

To validate the utility of iATPSnFR2 (and its affinity derivatives, A95K and A95K.S29W) for measuring changes in ATP production or consumption, we first tested in two related cell lines, LMTK-, a mouse fibroblast cell line, and a Rhoº (ρ^0^) derivative, which lacks mitochondrial DNA(15) observing changes in fluorescence in response to treatment with different ATP-affecting drugs. Because most cell lines are heavily dependent on glycolysis for energy production(16), we expected a significant decrease in fluorescence upon treatment with 2-deoxyglucose (2dOG). 2dOG can be phosphorylated by hexokinase, the first enzymatic step in glycolysis, but its product (2dOG-P) cannot be further processed, resulting in competitive inhibition of phosphoglucose isomerase(17) and therefore intracellular ATP depletion.

Prior to treatment, the green:red ratio of imaged cells was greater when the cells expressed higher affinity variants of iATPSnFR2, indicating that affinity affected ATP saturation. (A95K.S29W > A95K > A95A.A119L. Fig. 1). When glucose was replaced with 2dOG, green fluorescence rapidly dropped in the ρ^0^ cells (Fig. 1c), which have no alternative source of ATP other than glycolysis, with the weakest affinity variant (A95A.A119L) becoming nearly non-fluorescent within minutes of treatment, followed by the next highest affinity variant (A95K) and finally, the highest affinity variant (A95K.S29W). Meanwhile, 2dOG treatment resulted in decreased fluorescence only in the parental LMTK-cells expressing the weakest affinity variant (A95A.A119L), indicating that alternative methods for generating ATP, namely through mitochondrial oxidative phosphorylation, were available (Fig. 1d). When the parent cells were treated with both 2dOG and oligomycin (to inhibit oxidative phosphorylation), the fluorescence of the parent cells dropped in a manner similar to the ρ^0^ cells (Fig. 1e).

The self-consistent result of the three affinity variants responding to 2dOG as expected indicates that the primary effect of the treatment is due to decreasing [ATP], and not some other metabolic change, such as pH, [ADP] etc. To further corroborate that, we also used cpSFGFP (without an ATP binding subunit) as a “null” sensor. Since cpSFGFP and iATPSnFR2 have very similar pH dependencies, the former should provide an appropriate reporter for how treatment might affect intracellular pH or other artefactual changes in fluorescence. In ρ^0^ cells expressing either cpSFGFP or iATPSnFR2.A95A.A119L, replacement of buffer containing 10 mM glucose with fresh buffer (also containing 10 mM glucose), causes a transient increase in the green fluorescence, though the increase is greater for cells expressing iATPSnFR2 than cpSFGFP (Supp. Fig. 2), indicating that addition of fresh glucose probably gives the cells a small boost in ATP production. (Note: a transient apparent change in [ATP] upon treatment is also observed with the ChemoG-ATP_SiR_ sensor (Fig. 3e of ref(3)). When the buffer is replaced by 10 mM 2dOG and later rescued with 10 mM glucose (Supp. Fig. 2), only cells expressing iATPSnFR2 show a significant decrease in fluorescence, while those expressing cpSFGFP remain mostly constant, confirming that iATPSnFR2 is primarily reporting Δ[ATP], not ΔpH.

**Figure 2.**
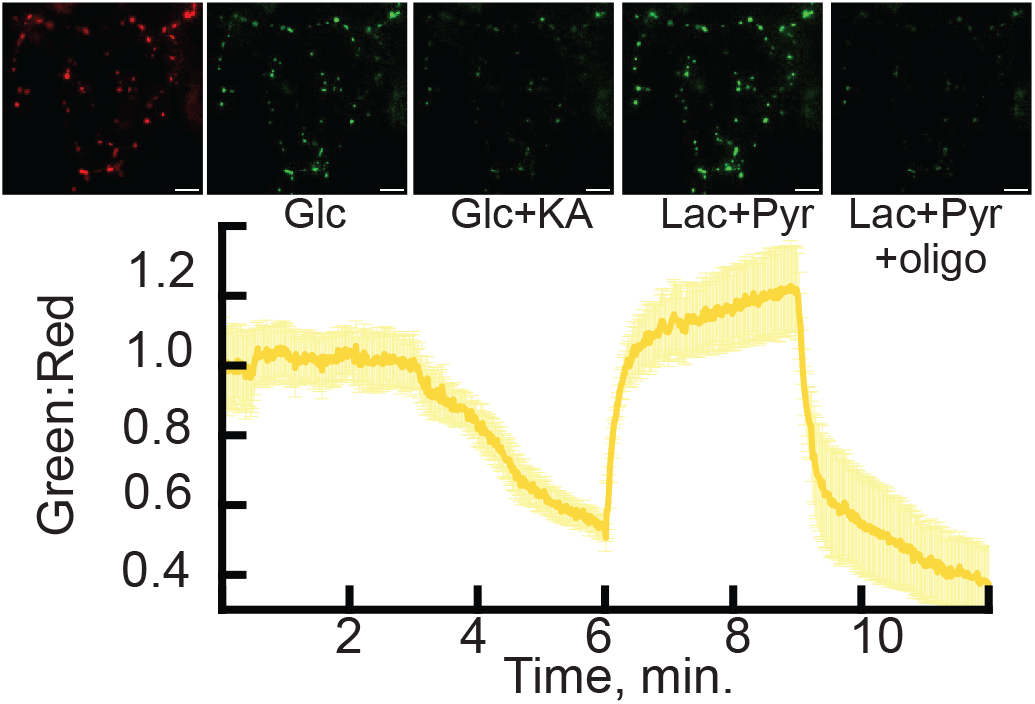
Response of iATPSnFR2 in matrix of axonal mitochondria to glycolytic and mitochondrial ATP synthase inhibitors. iATPSnFR2.A95A.A119L.HaloTag targeted to mitochondria with 4x-COX8 signal sequence was imaged for HaloTag-JF635 in 5 mM glucose. iATPSnFR2.A95A.A119L was imaged with perfusion of buffers containing: 5 mM glucose, 5 mM glucose + 10 μM koningic acid (KA), 1.25 mM lactate + 1.25 mM pyruvate, and finally 1.25 mM lactate + 1.25 mM pyruvate + 10 μM oligomycin. Scale bar: 10 μm. Data is represented as mean ± S.E. Images at top align with treatments and trace below. JF635 channel (red image) remained stable throughout the experiment.

**Figure 3.**
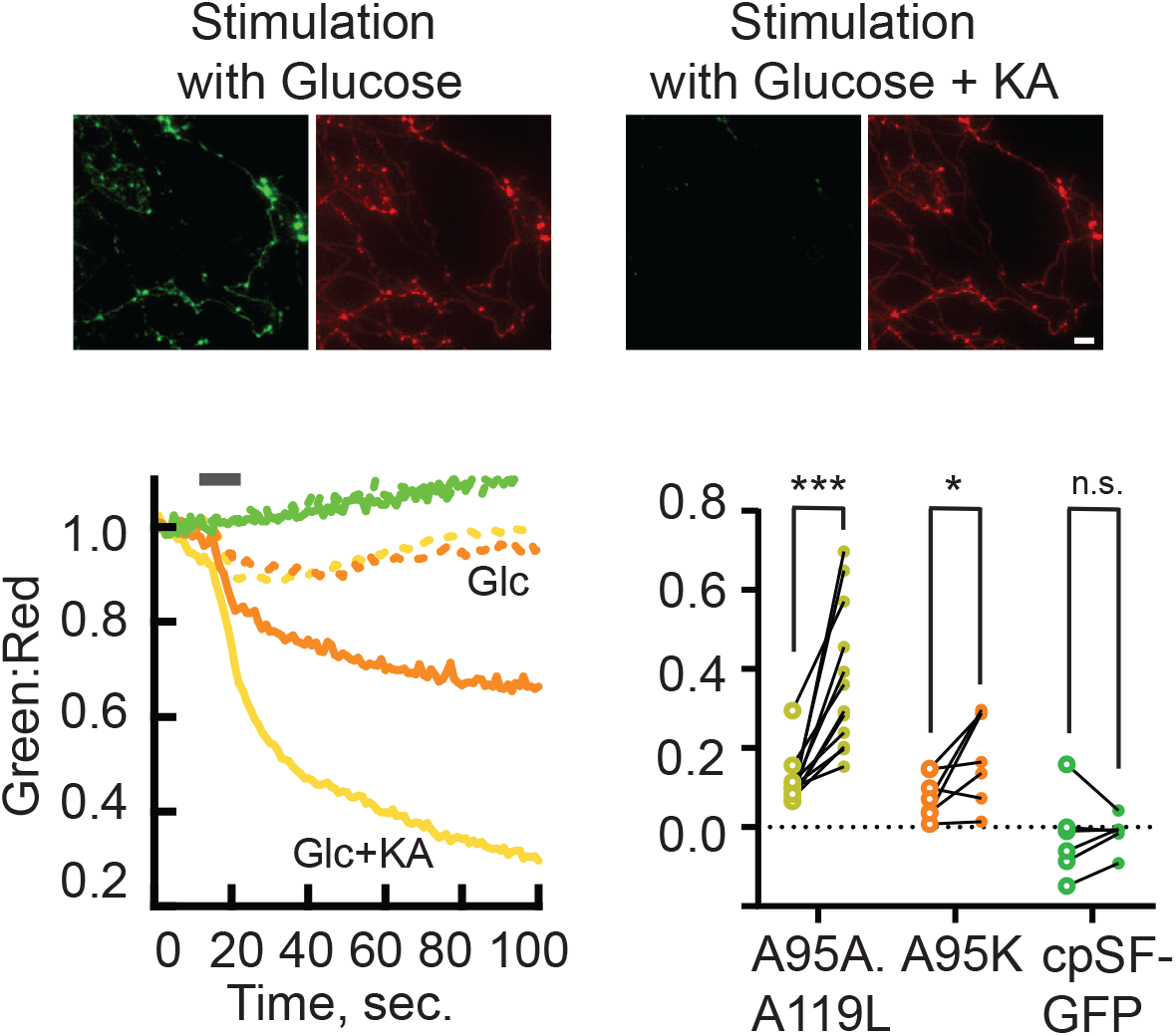
Depletion of cytosolic ATP in nerve terminals during action potential firing. Bottom left: Ratio of green to red fluorescence of axonal terminals in cultured neurons expressing iATPSnFR2.HaloTag-JFX650 during a burst of AP firing (6 sec at 50 Hz (grey bar)) in either 5 mM glucose (dashed lines) or 5 mM glucose + 10 μM koningic acid (solid lines). Low affinity variant A95A.A119L in yellow, medium affinity variant A95K in orange. Identical experiments carried out using cpSFGFP control (green) showed no change for the same stimulus for either condition. Top: images at the completion of the experiment in both green and red channels; scale bar 10 μm. Bottom Right: Maximum change in ratio during the stimulation period in the absence of KA (open circles) or presence of 10 μM KA (filled circles).

### Use of iATPSnFR2 to monitor consumption of ATP in primary neurons

Neurons are highly polarized cells that consist of soma, axons, and dendrites. The generation and consumption of ATP in these different sub-compartments are also semi-autonomously regulated. Genetically encoded sensors can be targeted to specific subcellular compartments by using different signal sequences and can be used to monitor [ATP] during metabolic perturbations. To that end, we fused four copies of the COX8 leader sequence(18) to the N-terminus of iATPSnFR2.A95A.A119L.HaloTag (mito-iATPSnFR2.HaloTag) to target it to the matrix of mitochondria. Expressing the targeted sensor in primary hippocampal neurons led to a distinct punctate appearance in both the red (here visualized with JF635 liganded to HaloTag) and green channels (Fig. 2), consistent with mitochondrial targeting. Application koningic acid (KA), a covalent inhibitor of glyceraldehyde 3-phosphate dehydrogenase (GAPDH) (19) blocked glycolysis and therefore, resulted in a gradual decrease of [ATP] (Fig. 2), while JF635 signal remained unperturbed (Supp. Fig. 3a). Bath application of a mixture of lactate and pyruvate fully restored the ATP signal, as expected since this allows it to bypass the glycolytic block, directly fueling the mitochondrial tricarboxylic acid cycle. Subsequently blocking the mitochondrial F_0_-F_1_ ATPase with oligomycin led to a further collapse of the ATP signal (Fig. 2). Similar results were obtained using mito-iATPSnFR2.mIRFP670nano3 (Supp. Fig. 3b).

One key question concerning the biology of energy consumption in neurons is the nature of the mechanisms that maintain the balance between ATP consumption and ATP production during electrical activity. While this has previously been addressed by our use of luciferase(10), the time resolution of that experiment was on the order of one minute. Here we used iATPSnFR2.A95A.A119L.HaloTag-JFX650 to observe ATP dynamics with sub-second time resolution, during a 6-second window of intense action potential firing at nerve terminals of dissociated hippocampal neurons in culture (Fig. 3). When we performed the stimulation under normal conditions, fluorescence dropped slightly, and recovered to baseline within one minute of concluding the stimulation. When we subsequently inhibited production of ATP by glycolysis with KA, we observed an even greater decrease in fluorescence that did not recover within the minute and a half observation window. The use of the higher affinity iATPSnFR2.A95K.HaloTag-JFX650 was still able to detect the activity changes in fluorescence signal, albeit with a much smaller amplitude. The negative control cpSFGFP (without an ATP-binding domain) remained unchanged even after ATP depletion. These data recapitulate our previous observations that nerve terminals have a robust on-demand ATP synthesis program that responds rapidly to electrical activity (10), while improving the time resolution by an order of magnitude.

### iATPSnFR2 reveals differences in ATP consumption among individual boutons

We previously showed that, even in the absence of electrical activity, nerve terminals have high basal ATP consumption but both single bouton and kinetic details were limited by the low photon flux of the luciferase-based reporter(20). We repeated these experiments using synaptically targeted iATPSnFR2 that was fused to synaptophysin(21), sending it to the synaptic vesicle surface, facing the cytosol. Similar to previous findings, when ATP production was blocked (in this case by inhibiting GAPDH with KA) in the presence of the Na^+^ channel blocker tetrodotoxin (TTX), the average green to red fluorescence ratio of iATPSnFR2.HaloTag-JFX650 (averaged over 7 different neurons, each contributing 20-30 boutons) gradually declined over a 5-10 min period as previously reported(20). The weaker A95A.A119L variant showed a greater drop than the medium affinity A95K variant (Fig. 4a). In contrast to what we observed in the mitochondrially-targeted sensor, when using JF635 as a HaloTag ligand, the red channel fluorescence was not inert with respect to ATP changes (Supp. Fig. 4a). Rhodamine-based dyes can interact with ATP on their own(22), and it appears that HaloTag-JF635 itself is sensitive to ATP, albeit with low ΔF/F (Supp. Fig. 4b). This confounding artefact was minimized by using a newer generation HaloTag dye, JFX650 (Supp. Fig. 4a). The improved signal to noise properties of iATPSnFR2 allowed us to extract several new features regarding single synapse ATP control. Comparisons of the initial ratio values across synapses and experiments showed that across boutons, the resting ATP value varied as much as 4-fold, and by 50% across cells, but this variation was minimized following ATP depletion by KA (Fig. 4b). Similarly, the kinetics of ATP depletion in individual nerve terminals behaved unexpectedly. Although the average behavior across all boutons shows a gradual decline in ATP (Fig. 4a), no individual bouton looked like the average, displaying instead two distinct kinetic features that varied widely across boutons (Fig. 4c). At the single bouton level, ATP signals declined very rapidly (typically showing depletion within ∼ 10-20 sec) but only after a delay time that varied significantly across boutons. Individual bouton ATP levels under these conditions were best described by a simple Boltzmann function, 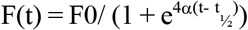 where t_½_ describes the delay time to the precipitous drop in signal at a rate α (see methods) (Fig. 5d). Both these kinetic parameters, t_½_ and α varied significantly across the population of nerve terminals. A correlation analysis of the two extracted parameters showed that in general smaller t_½_ were associated with higher values of α (Fig. 4e) and vice versa. Some cells showed tighter clustering of rate and t_½_ parameters than others (Fig. 4f).

**Figure 4.**
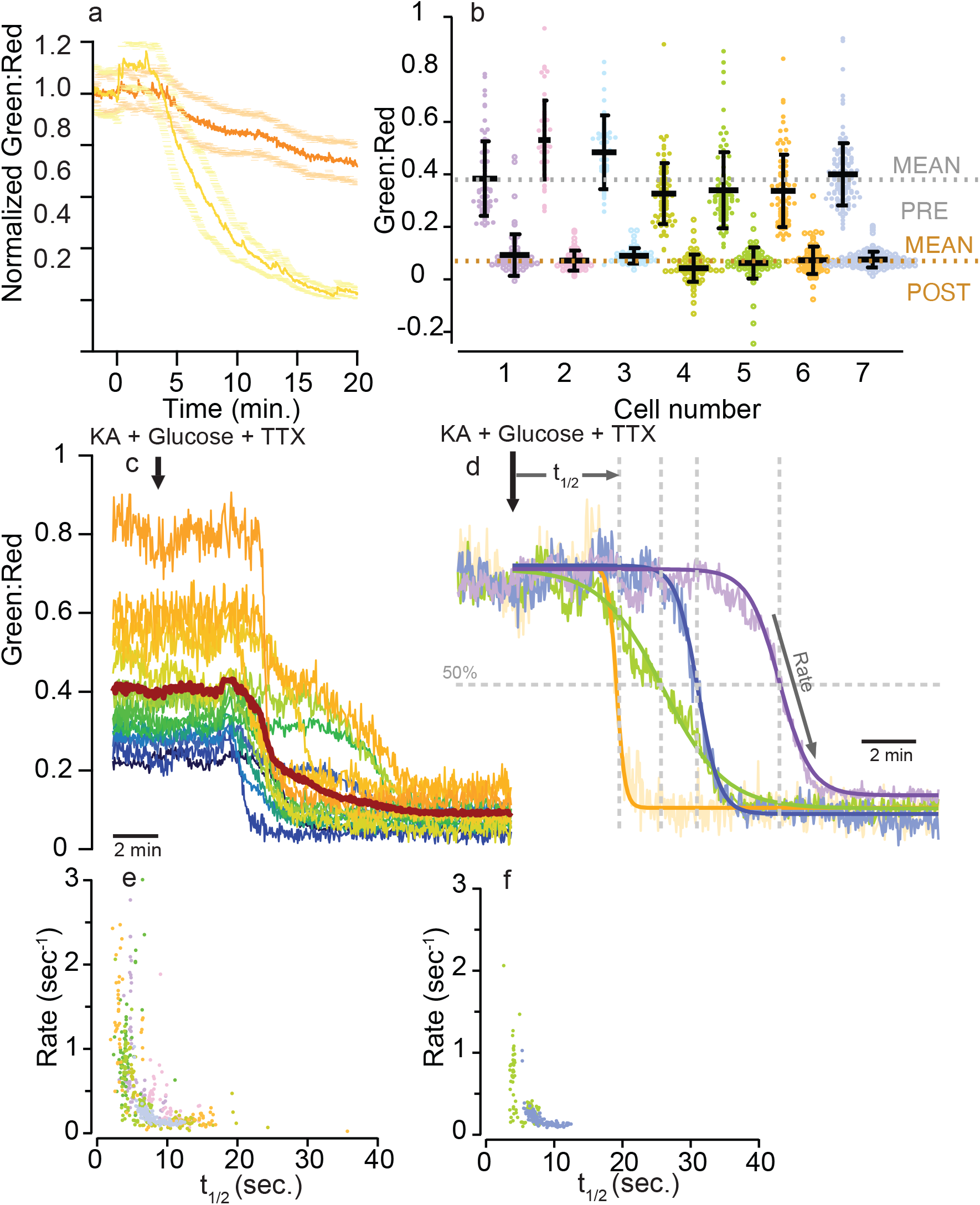
Spontaneous depletion of ATP in boutons detected by iATPSnFR2. iATPSnFR2 was targeted to boutons by fusing it to the C-terminal of synaptophysin. a) Average ratio of green to red fluorescence of iATPSnFR2-HaloTag-JFX650 normalized to pre-treatment baseline. Yellow, low affinity variant A95A.A119L (n=7); orange, medium affinity variant A95K (n=6), ± s.e. Treatment with 5 mM glucose + 10 μM KA + 0.3 μM TTX initiated at t = 0. b) Variation in ATP across individual boutons and across individual neurons pre- and post-ATP depletion by koningic acid of the 7 cells shown in a) with the low affinity variant. c) Representative raw traces from individual boutons expressing A95A.A119L variant. d) Normalization of representative traces illustrate that individual boutons have unique half-times (t_½_) to ATP depletion and unique rates of ATP consumption. e) Cross correlation of single bouton depletion rates versus t_½_ values shows there is in general an inverse relationship between these parameters, with some cells occupying regions of the parameter space. f) Boutons from only two cells (cells 4 & 7 in b) shown for clarity.

## Discussion

Here we present an improved ATP sensor -iATPSnFR2, optimized for both higher fluorescence changes in response to saturating ATP (ΔF/F ∼ 12), and modulation of its affinity for ATP such that its range of maximal sensitivity (*K*_d_) matches the expected range of ATP concentrations within normally functioning cells, including neurons. We also provide two higher affinity variants to serve as controls or for use in other cell types or cellular sub-compartments that might be expected to have lower concentrations of ATP. We validate its utility for measuring changes in cytosolic ATP using cellular imaging in cultured cell lines, perturbed with different pharmacological compounds known to affect ATP production and consumption. Inclusion of a far-red fluorophore (*via* HaloTag or mIRFP670nano3) allows for ratiometric measurement to account for expression and movement artefacts. iATPSnFR2 can be targeted to the axonal mitochondria, where it reports homeostasis of mitochondrial ATP in presence of a suitable fuel for the tri-carboxylic acid cycle. It also reports depletion and subsequent rescue of mitochondrial ATP after perfusion of oxphos substrate and inhibitors. Finally, by targeting it to boutons, we can observe spontaneous depletion of ATP and observe that each bouton has its own unique ATP consumption profile.

While multiple other fluorescence-based sensors have been developed, none of them have been widely adopted for determination of ATP consumption in cell culture or *in vivo*. ATP production pathways through glycolysis produces lactate and acidifies the extracellular space. ATP production from the electron transport chain in mitochondria consumes O_2_. The extracellular acidification rate (ECAR) and oxygen consumption rate (OCR) can be measured and used as indirect proxies for ATP consumption and are the underlying technologies of the Seahorse analyzer(23). Typically, these rates are measured in cell culture under normal/healthy conditions to establish a baseline, and then mitochondrial inhibitors (oligomycin and rotenone/antimycin A, but sometimes other inhibitors) are added to drop mitochondrial ATP production to zero, and through a series of complex calculations, ATP production rates, divided between mitochondrial or glycolytic production, are determined.

Measuring pH and O_2_ consumption in cell culture is useful for determining the energetic expenditure differences between cell types or between cell cultures that have been treated with different drugs, but the technique also has significant limitations. Primarily, measuring pH changes and O_2_ consumption are just proxies for ATP production, and not a direct measurement. Other metabolic events or perturbations that affect pH and O_2_ use will be interpreted as ATP consumption. Importantly. The Seahorse system is limited to cell culture analysis and is not compatible with *in vivo* applications or measurement of ATP consumption with other manipulations, such as electrical stimulation. This technology also affords no spatial resolution either at the single cell or subcellular length scale. Furthermore, once cells have been treated with mitochondrial poisons, they are no longer viable and cannot be assayed again. Finally, the time course of measurement using such devices requires integration of signals over time and is not compatible with sub-second resolution. We expect that iATPSnFR2 will provide researchers with previously exciting new opportunities to study ATP dynamics with temporal and spatial resolution that has, until now, been unavailable.

## Materials and Methods

### Mutagenesis and screening of bacterial lysates

iATPSnFR1 was cloned into a bacterial expression vector derived from pRSET (Invitrogen) with restriction sites designed to make downstream subcloning into mammalian expression vectors easier. Site saturation mutagenesis was performed using the uracil template method(24), as described previously(25). Mutagenesis reactions were transformed into T7 express bacterial cells (New England Biolabs) and plated on LB-Amp agar plates. Individual colonies were picked into 2 mL, square well, 96-well plates filled with 0.9 mL auto induction media(26) and grown overnight at 30°C with rapid shaking (350+ RPM). To remove excess ATP, 96-well plates were centrifuged, pellets resuspended in Tris Buffered Saline, and re-centrifuged, a total of 3 times, then frozen overnight as dry pellets. iATPSnFR variants were assayed in bacterial lysate by addition of 0.9 mL Mammalian Cell Imaging Buffer (20 mM HEPES, 119 mM NaCl, 2.5 mM KCl, 2 mM MgCl_2_, 2 mM CaCl_2_, pH 7.2), rapid vortexing, then pelleting by centrifugation. Clarified lysate was transferred to a black 96-well plate (Greiner). Fluorescence was measured in a plate reader (Tecan Spark) with 485 nm excitation (20 nm bandpass) and 535 nm emission (20 nm bandpass). ATP (Sigma) was added to 1 μM and fluorescence re-measured. ATP was added to 1 mM and fluorescence re-measured. Variants with responses to either concentration with high ΔF/F were carried forward with mutations of “linker 1” until one of those winners, L1-VLVG was good enough to become the template for mutations at “linker 2”.

### Protein expression and purification and characterization

iATPSnFR2 variants were transformed into E. coli BL21(DE3) pLysS cells. Protein expression was induced by growth in 300 mL autoinduction media(26) supplemented with 100 μg/mL ampicillin at 30°C. Proteins were purified by immobilized Ni-NTA affinity chromatography(27). iATPSnFR proteins were eluted with a 120-mL gradient from 0 to 200 mM imidazole. Fractions that were fluorescent were pooled and concentrated by ultrafiltration (Amicon) and then dialyzed by Slide-A-Lyzer cassette (30 kDa cutoff) in Mammalian Cell Imaging Buffer. Protein concentration was quantified by alkali denaturation and measurement of the GFP chromophore, with an extinction coefficient of 44,000 M^-1^ cm^-1^ at 447 nm.

### Synthetic fluorophore conjugation

Purified protein was mixed with 1.1 molar equivalents of JFX650 HaloTag Ligand and incubated at room temperature for 1 hour and then at 4°C overnight. Excess HaloTag Ligand was removed by gel filtration on a PD-10 column (GE Life Sciences) and re-concentrated by ultrafiltration.

### Equilibrium measurements

All *in vitro* assays were performed in Mammalian Cell Imaging Buffer unless otherwise noted. iATPSnFR2 equilibrium measurements (affinity, specificity, pH) were performed with 0.2 μM protein in a Tecan Spark fluorimeter with 20 nm bandpass windows and excitation at 485 nm and emission at 535 nm.

### Kinetic measurements

iATPSnFR2 protein (0.2 μM) was rapidly mixed with an equal volume of stock ATP solution in an Applied Photophysics SX-20 stopped flow fluorimeter with a 490 nm LED excitation and 510 nm long pass filter for emission. Final concentration of protein was thus 0.1 μM. Concentrations of ATP listed in figures are the final concentration after mixing.

### Cloning into mammalian expression vectors

iATPSnFR2.HaloTag variants were subcloned by restriction digest into an AAV vector with a CAG promoter *via* BglII/PstI digest from the bacterial expression vector. The pAAV.CAG vector includes an extra serine after the initial methionine and lacks a polyhistidine tag. To target the sensor to the mitochondrial matrix, four repeats of COX8(mito) leader sequence (SVLTPLLLRGLTGSARRLPVPRAK) separated by spacer (IHSLPPEGPW) was introduced at N-terminal of iAPTSnFR2(A95A/A119L). HaloTag was incorporated at the N-terminal of iATPSnFR2.A95A.A119L separated by a linker (LQSTGSGNAVGQDTQER). The plasmid was transformed into stbl3 competent *E*.*Coli* and DNA was harvested from 200 mL growth media using an endotoxin free plasmid purification kit (Qiagen).

### Rhoº cell culture

Rhoº and the parental LMTK-cells were maintained in T-75 flasks with DMEM + 10% FBS + 1 mM pyruvate + 5 μM uridine. Two days prior to transfection, they were split into a 24-well plate (half Rhoº, half LMTK) at 0.6 and 0.1 million cells per well, respectively. Cells were transfected with iATPSnFR2.HaloTag variants by lipofectamine (0.5 μg DNA + 2 μL lipofectamine 2000 per well) for 4-6 hours in DMEM lacking FBS or antibiotics. Transfection media was replaced with fresh DMEM + 10% FBS + penicillin/streptomycin overnight. Cells were labeled with JFX650 HaloTag ligand by adding the fluorophore to 100 nM for an hour. Cells were washed with Mammalian Cell Imaging Buffer and imaged in a Cytation 5 imaging reader (BioTek) over the course of about 2 hours. After about 20 minutes of imaging to establish a baseline fluorescence, the culture plate was removed from the instrument and buffer was replaced with the query buffer. The equilibration buffer contained 10 mM glucose, and the query buffers contained 10 mM 2dOG.

### Rhoº cell image processing

Images were background subtracted and aligned for movement artefacts with the Gen5 software package that controls the Cytation 5 instrument. Images were assembled as TIF stacks and imported into ImageJ. For each well, an automated, custom script identified regions of interest (ROIs) by thresholding on the far-red channel and expanding by 2 pixels. Those ROIs were used to determine the ratio of green to red fluorescence for each well. ROIs were curated to remove those that outlined cells that detached or appeared to become spherical during the experiment.

### Animals

All experiments involving animals were performed in accordance with protocols approved by the Weill Cornell Medicine Institutional Animal Care and Use Committee. Neurons were derived from Sprague-Dawley rats (Charles River Laboratories strain code: 001, RRID: RGD_734476) of either sex on postnatal days 0–2).

### Neuronal cell culture

Primary neuronal cultures were prepared as previously described(28). Hippocampal CA1 to CA3 regions were dissected, dissociated, and plated onto poly-L-ornithine-coated coverslips. Plating media consisted of the minimal essential medium, 0.5% glucose, insulin (0.024 g/l), transferrin (0.1 μg/l), GlutaMAX 1%, N-21 (2%), and fetal bovine serum (10%). After 1–3 days *in vitro* (DIV), cells were fed and maintained in media with the following modifications: cytosine β-D-arabinofuranoside (4 μM) and FBS 5%. Cultures were incubated at 37 °C in a 95% air/5% CO2 incubator. Calcium phosphate-mediated gene transfer was performed on DIV 6–8, and neurons were used for experiments on DIV 14–21.

### Neuronal cell imaging

Coverslips were loaded onto a custom chamber, mounted on a Zeiss inverted microscope and perfused at ∼100 μl min^−1^ with Tyrode’s solution containing: 119 mM NaCl, 2.5 mM KCl, 30 mM glucose, 25 mM HEPES, 2 mM CaCl_2_, 2 mM MgCl_2_, 50 μM DL-2-amino-5-phosphonovaleric acid (APV, AP5), 10 μM 6-cyano-7-nitroquinoxaline-2,3-dione (CNQX), adjusted to pH 7.4. Prior to each experiment, a JF-dye aliquot (JFX650 or JF635) was diluted (100 nM final) into the cell culture medium, incubated for 20 min and washed thoroughly twice (5 min each) with imaging buffer. The temperature was maintained at 37°C with a custom-built objective heater under feedback control (Minco). Fluorescence was stimulated with OBIS 488 nm LX or OBIS 637 nm LS lasers (Coherent) passing through a laser speckle reducer (LSR 3005 at 12° diffusion angle, Optotune). Emission was acquired with a 40x, 1.3 numerical aperture oil immersion objective (Fluar, Zeiss) on an Andor iXon + Ultra 897 electron-multiplying charge-coupled device camera. Action potentials (APs) were evoked with platinum-iridium electrodes generating 1 msec pulses of an electric field of 10 V cm^−1^ via a current stimulus isolator (A385, World Precision Instruments). Laser power at the back aperture was ∼0.32 mW for 637 nm and ∼0.52 mW for 488 nm. Laser wavelength excitation was alternated between frames during image acquisition, with an exposition of 100 msec at 1 Hz acquisition frequency, A custom-designed Arduino board coordinated AP and laser stimulation with frame acquisition.

### Synaptic Bouton and axonal mitochondria Image analysis

Time series of imaging pairs (HaloTag and iATPSnFR2) were automated split into two independent image series using a custom-written Fiji routine to facilitate analysis. ATP signals are reported as a ratio between iATPSnFR2:HaloTag (Green:Red). Images were analyzed using the ImageJ plug-in Time Series Analyzer V3 where 50 to 100 circular regions of interest (ROIs) of radius 1 um corresponding to synaptic boutons or mitochondria expressing the Syn-iATPSnFR2.HaloTag or mito-iATPSnFR2.Halo (Halo dye positive) were selected. Image loading and posterior raw data saving were automatized using a homemade Python code for Fiji. Synaptic boutons signals were analyzed using homemade script routines in Igor-pro v6.3.7.2 (Wavemetric, Lake Oswego, OR, USA). ATP ratio signal (Green:Red) was calculated per individual bouton, normalized to the baseline and fitted with a Boltzmann’s sigmoidal function from the time that 10uM koningic acid was administered, as follows:

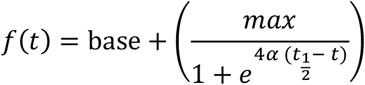

Where t_1/2_ is the t value where Y is at (base + max)/2, and α represents the rate of signal drop derived from:

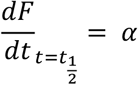

For electrical activity experiments, single cell data was filtered by excluding any cell data where no ATP change was observed when cells were stimulated in presence of KA.

## Acknowledgments

We would like to thank Anastasia Osowski (Janelia) for help in formatting the manuscript. This research was supported in part by NIH grants (NS036942 & NS11739) to TAR and in part by Aligning Science Across Parkinson’s ASAP-000580 through the Michael J. Fox Foundation for Parkinson’s Research (MJFF). For the purpose of open access, the authors have applied a CC BY public copyright license to all Author Accepted Manuscripts arising from this submission.

