## Supplementary material for "iATPSnFR2: a high dynamic range fluorescent sensor for monitoring intracellular ATP": Supp Figs

**Supplementary Figures**

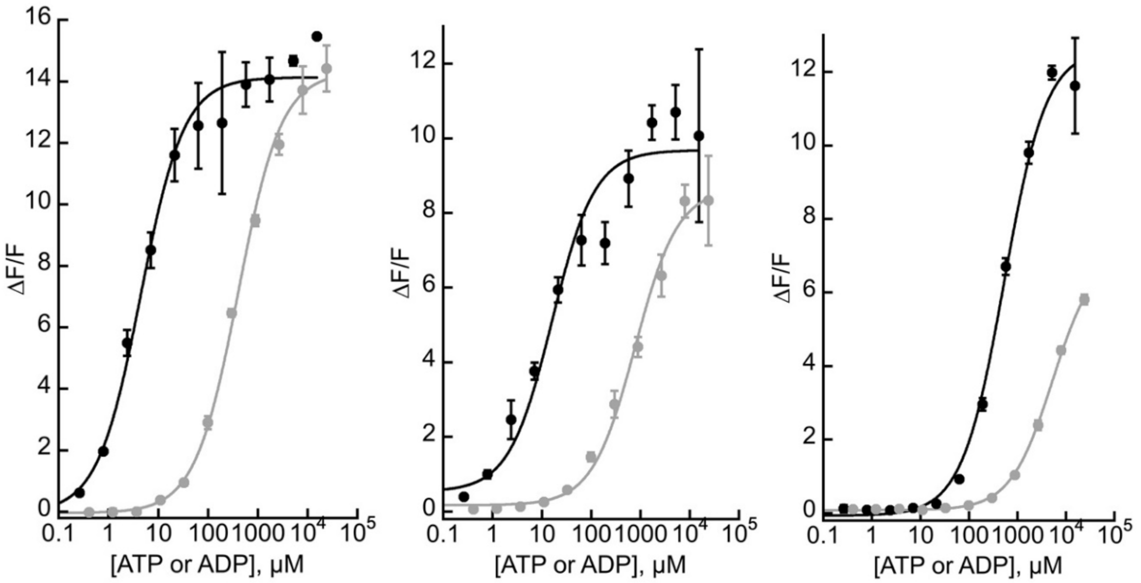


**Supplementary Figure 1a.** <u>*In vitro* characterization.</u>
iATPSnFR2 affinity for ATP and decoys. Three variants of iATPSnFR2.HaloTag-JFX650 titrated with ATP (black) and ADP (grey), head-to-head. S29W.A95K (left); A95K (center); A95A.A119L (right). Binding curve fits, in μM, for ATP and ADP are S29W.A95K: 4 μM, 400 μM. A95K: 16 μM, 780 μM. A95A.A119L: 530 μM, 5300 μM.

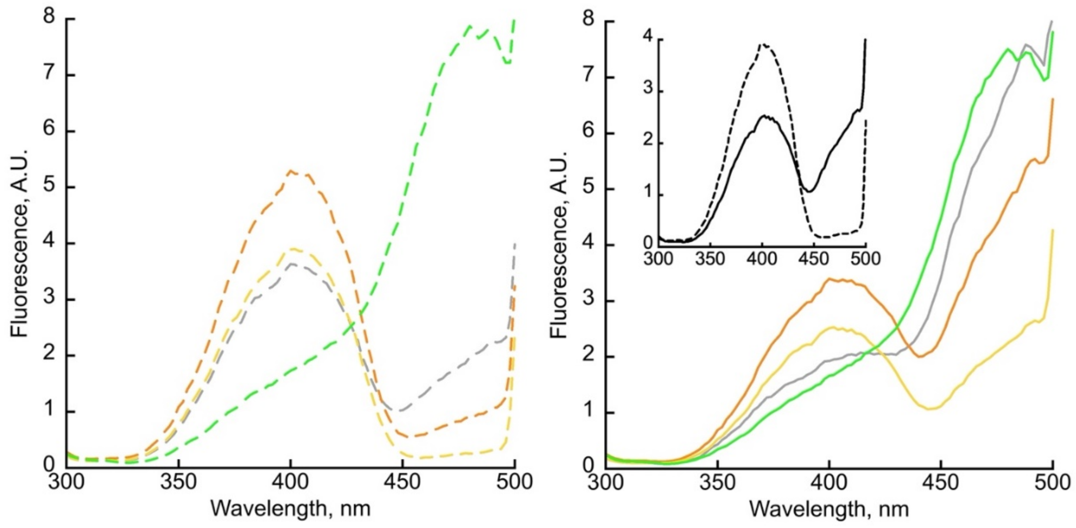


**Supplementary Figure 1b.** *In vitro* characterization.

Excitation spectra of iATPSnFR2.S29W (grey), .A95K (orange), .A95A.A119L (yellow), and cpSFGFP (green) in the absence of ATP (left, dashed lines) and in the presence of 7 mM ATP. Emission observed at 515 nm (5 nm bandpass). Excitation scanned with 5 nm bandpass. Inset shows A95A.A119L variant spectra ± ATP to indicate the isosbestic point.

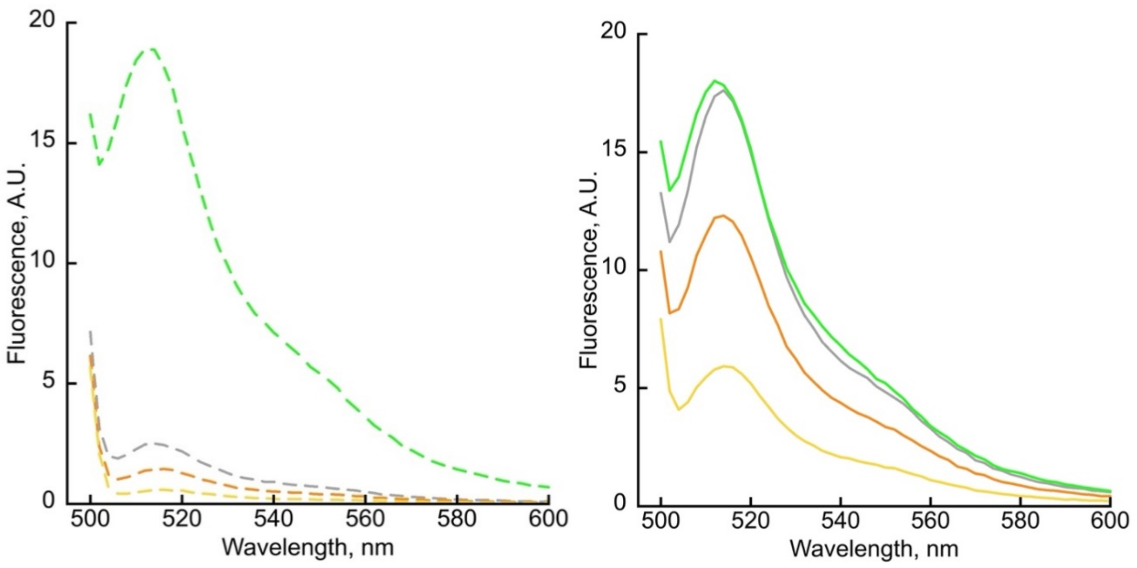


**Supplementary Figure 1c.** *In vitro* characterization.

Emission spectra of iATPSnFR2.S29W (grey), .A95K (orange), .A95A.A119L (yellow), and cpSFGFP (green) in the absence of ATP (left, dashed lines) and in the presence of 7 mM ATP. Excitation at 485 nm (5 nm bandpass). Emission scanned with 5 nm bandpass.

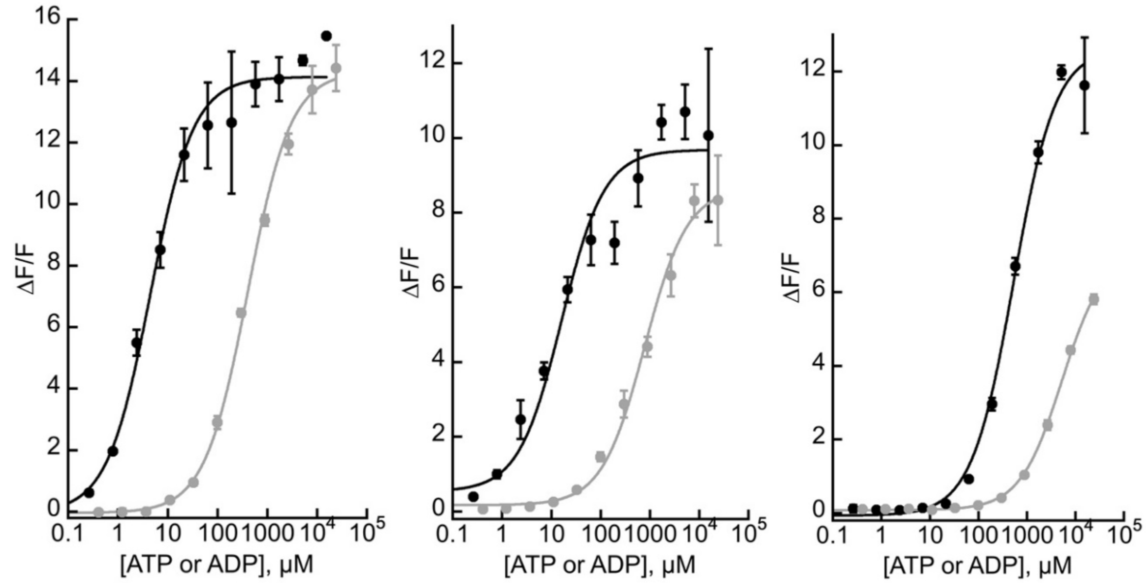


**Supplementary Figure 1d.** _In vitro_ characterization.
iATPSnFR2 affinity for ATP and decoys. Three variants of iATPSnFR2.HaloTag-JFX650 titrated with ATP (black) and ADP (grey), head-to-head. S29W.A95K (left); A95K (center); A95A.A119L (right). Binding curve fits, in μM, for ATP and ADP are S29W.A95K: 4 μM, 400 μM. A95K: 16 μM, 780 μM. A95A.A119L: 530 μM, 5300 μM.

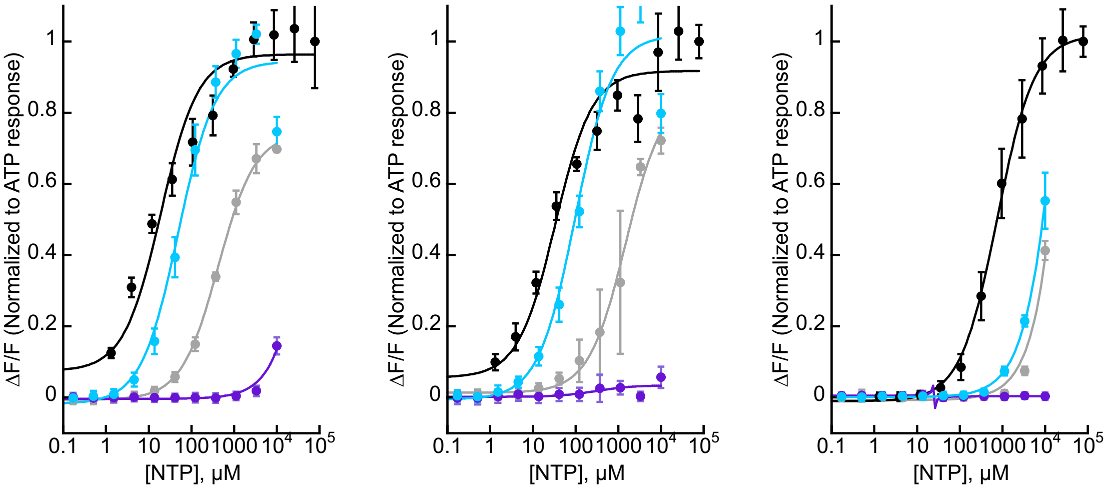


**Supplementary Figure 1e.** *In vitro* characterization.

iATPSnFR2 affinity for ATP and other nucleoside triphosphates. Three variants of iATPSnFR2.HaloTag-JFX650 titrated with ATP and other NTPs, head-to-head. S29W.A95K (left); A95K (center); A95A.A119L (right). ATP (black), CTP (cyan), GTP (grey), TTP (purple). Binding curve fits, in µM, for ATP, CTP, GTP are S29W.A95K: 20 µM, 50 µM, 430 µM. A95K: 30 µM, 100 µM, 1500 µM. A95A.A119L: 750 µM, ~10 mM, ~10 mM.

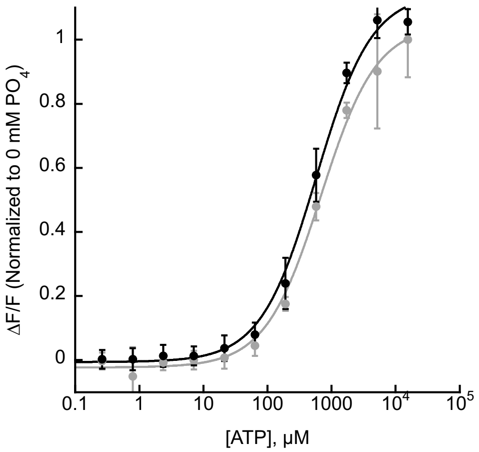


**Supplementary Figure 1f.** *In vitro* characterization.
Decoys. The A95A.A119L.HaloTag-JFX650 sensor is negligibly affected by inorganic phosphate. Head-to-head titration of ATP without added phosphate (black) and with 20 mM PO4 (grey) (diluted from 1 M NaPO4 stock, pH 7).

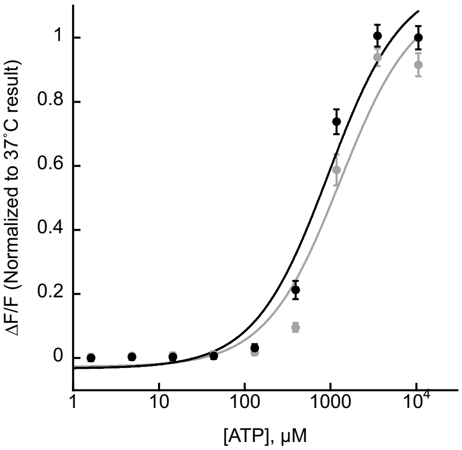


**Supplementary Figure 1g.** *In vitro* characterization.
Temperature dependence. ATP titration of iATPSnFR2.A95A.A119L variant at 25˚C (grey) and 37˚C (black). Error bars are s.d. of three technical replicates. Fluorescence was measured in a Cytation 5 plate reader, which has relatively rapid heating.

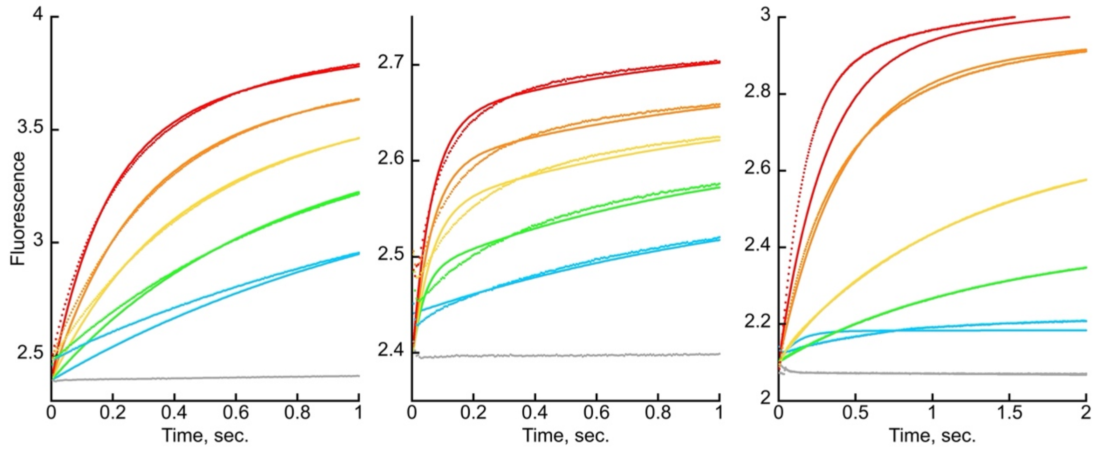


**Supplementary Figure 1h.** *In vitro* characterization.
Kinetics. The three different variants of iATPSnFR2 bind ATP within a one or two seconds of rapid mixing. S29W.A95K (left) and A95K (center) bind ATP faster than A95A.A119L (right). The data from stopped flow fluorescence do not fit to a single exponential (not shown), but very well to the addition of two exponentials (shown), indicating a more complicated, three-state mechanism of binding and fluorescence change. Panels are the sensor with a C-terminal fusion to HaloTag-JFX650. Final concentrations of ATP are: 10000, 3333, 1111, 370, 123 µM. Fluorescence data collected on an Applied Photophysics SX-20 stopped-flow apparatus with 490 nm LED excitation and 525 nm long pass filter. Data points are average of 5 technical replicates.

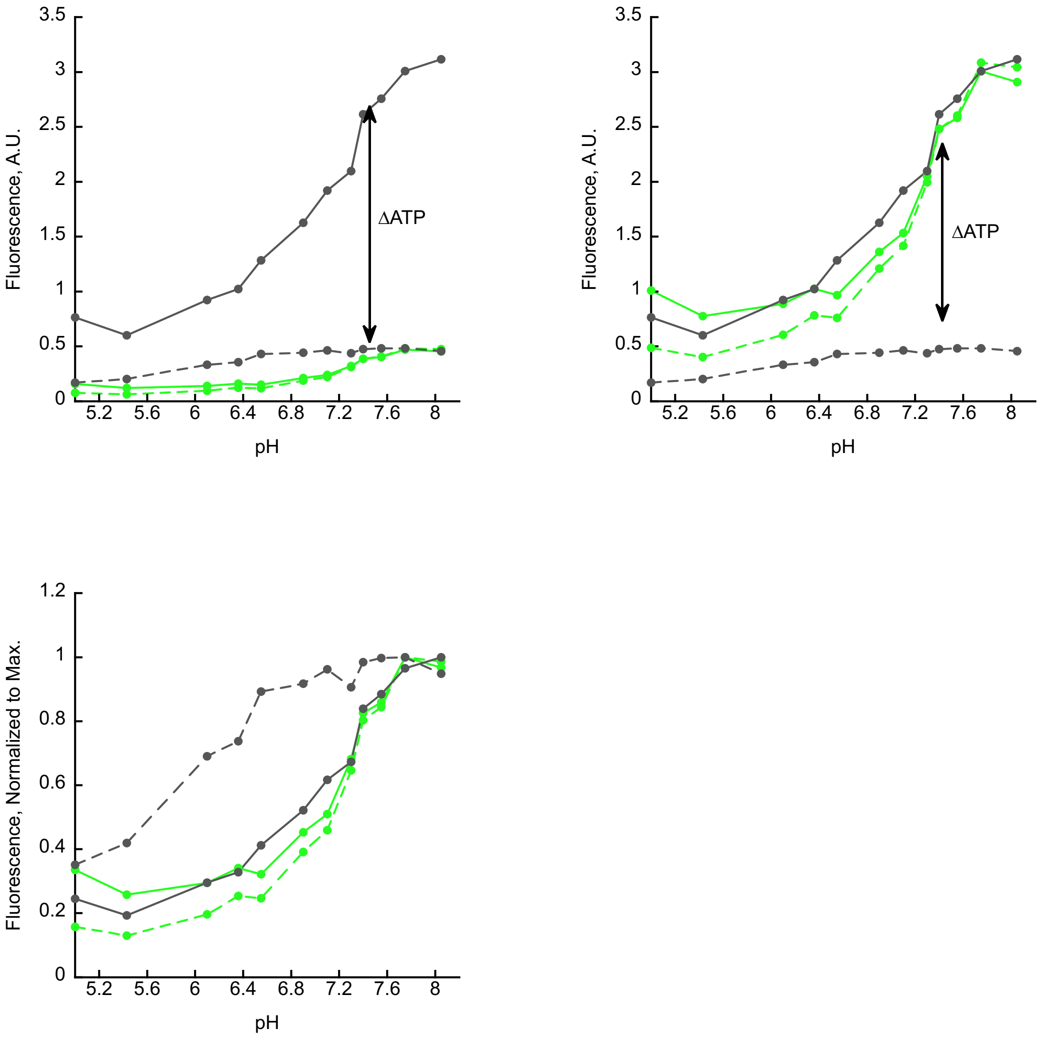


**Supplementary Figure 1i.** *In vitro* characterization.
pH dependence. iATPSnFR2.A95A.A119L.HaloTag and cpSFGFP were diluted to 0.2 µM in Mammalian Cell Imaging Buffer at variable pH. Fluorescence was measured at Ex/Em 485 nm / 535 nm (20 nm bandpass) or Ex/Em 600 nm / 630 nm (10 nm bandpass) without ATP (dashed lines). Then ATP was added to 2.5 mM from a concentrated stock and fluorescence was remeasured (solid lines). Each plot is a different presentation of the same data. In the top left panel, the fluorescence of cpSFGFP has been adjusted to approximately match that of ligand-free state of the sensor at pH 7.4. In the top right panel, the fluorescence of cpSFGFP has been adjusted to match that of the ATP-bound state of the sensor at pH 7.4. In the lower left plot, each data set has been adjusted so that its maximum value is 1. (Black, iATPSnFR2 green channel; green, cpSFGFP green signal).

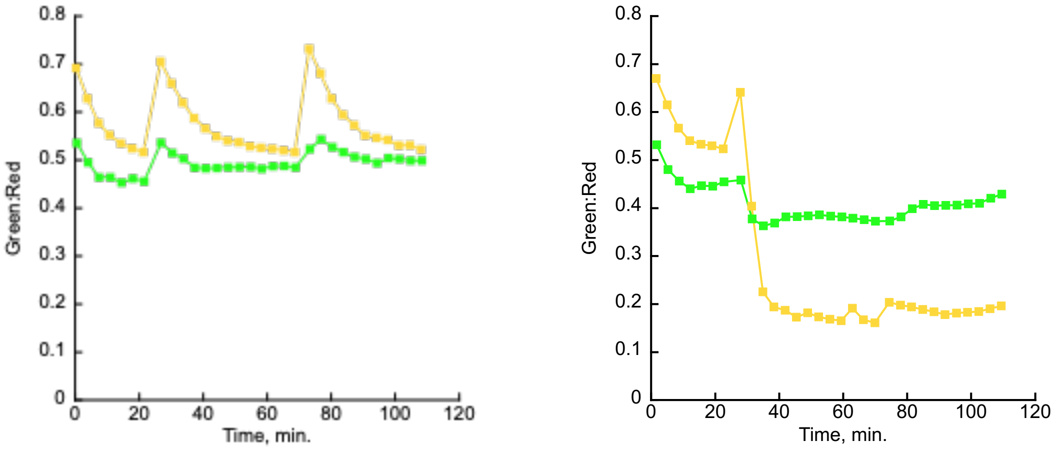


**Supplementary Figure 2.** Validation in fibroblast cell culture.
Left: Ratio of green to red (HaloTag-JFX650) fluorescence in $\rho^0$ cells transfected with iATPSnFR2.A95A.A119L variant (yellow) or cpSFGFP (green). Buffer containing 10 mM glucose was refreshed twice (at the points where fluorescence increased). Right: Same as on left, but with a switch from 10 mM glucose to 10 mM 2-deoxyglucose, and then back to 10 mM glucose.

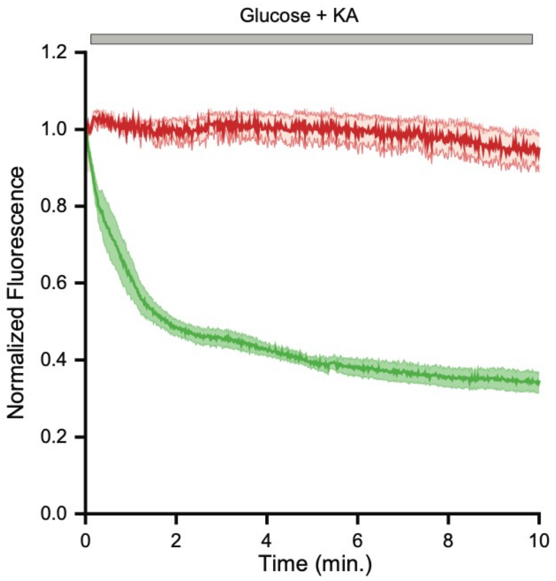


**Supplementary Figure 3a.** Validation of mito-iATPSnFR2.HaloTag in neuronal culture.
Fluorescence traces of both green (iATPSnFR2) and red (HaloTag-JF635) channels in neurons expressing mito-iATPSnFR2.A95A.A119L.HaloTag in mitochondria during perfusion with glucose and KA. Shown is the average fluorescence traces from 4 neurons (+/- SEM), with each neurons proving signals form 40-50 axonal mitochondria.

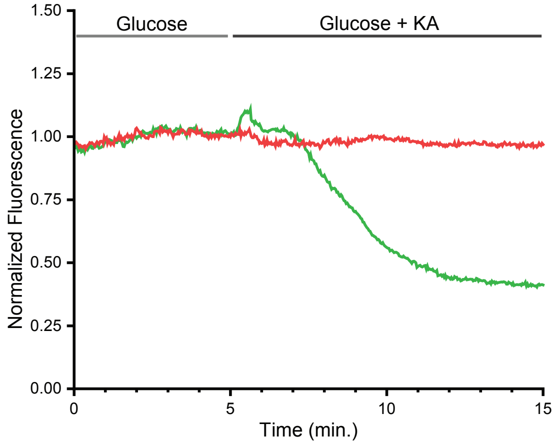


**Supplementary Figure 3b.** Validation of mito-iATPSnFR2.miRFP670nano3 in neuronal culture.
Average fluorescence traces (n=4) of both green (iATPSnFR2) and red (mIRFP670nano3) channels neurons expressing mito-iATPSnFR2.A95A.A119L.mIRFP670nano3 during perfusion with glucose and glucose with KA.

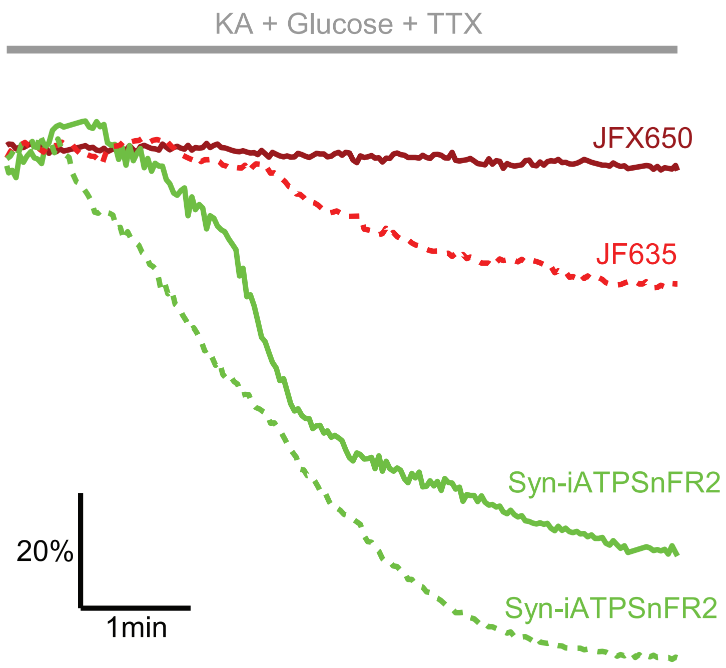


**Supplementary Figure 4a.** Validation of Syn-iATPSnFR2.HaloTag in axonal boutons.
Fluorescence traces of both green (iATPSnFR2 and red (HaloTag-JF635 or HaloTag-JFX650) channels in neurons expressing syn-iATPSnFR2.A95A.A119L.HaloTag on the cytosol-facing surface of synaptically targeted vesicles during perfusion with glucose and KA.

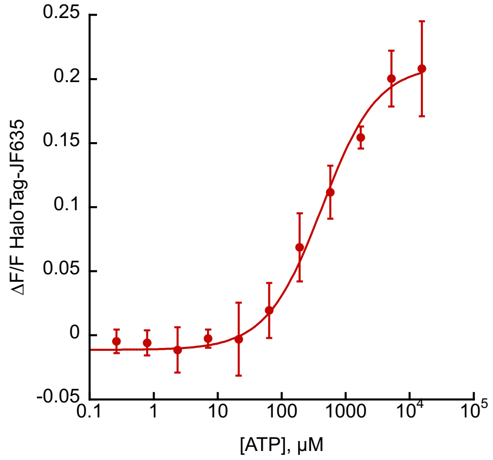


**Supplementary Figure 4b.** *In vitro* titration of HaloTag-JF635 with ATP.
Fluorescence of purified HaloTag-JF635 (0.2 uM in Mammalian Cell Imaging Buffer) was measured (Ex 625 nm, Em 670 nm, 20 nm bandpass) with varying concentration of ATP.
